# *Myoviridae* Phage *PDX* Kills Enteroaggregative *Escherichia coli* without Human Microbiome Dysbiosis

**DOI:** 10.1101/385104

**Authors:** Leah C. S. Cepko, Eliotte E. Garling, Madeline J. Dinsdale, William P. Scott, Loralee Bandy, Tim Nice, Joshua Faber-Hammond, Jay L. Mellies

**Affiliations:** 320 Longwood Avenue, Enders Building, Department of Infectious Disease, Boston Children’s Hospital, Harvard Medical School, Boston, MA 02115. U.S.A; Fred Hutchinson Cancer Research Center, 1100 Fairview Ave N, Seattle, WA, 98109. U.S.A; Biology Department, Reed College, 3203 SE Woodstock Blvd., Portland, OR, 97202. U. S. A; Department of Molecular Microbiology & Immunology, Oregon Health & Science University, 3181 SW Sam Jackson Park Road, Portland, OR 97239

**Keywords:** bacteriophage (phage), phage therapy, EAEC, Caudovirales, MDR, Myoviridae, Escherichia virus, microbiome, dysbiosis antibiotic alternatives.

## Abstract

**Purpose:** To identify therapeutic a bacteriophage that kills diarrheagenic enteroaggregative *Escherichia coli* (EAEC) while leaving the human microbiome intact.

**Methodology:** Phages from wastewater in Portland, OR, were screened for bacteriolytic activity using an overlay assay, and isolated in a sequential procedure to enrich for the recognition of core bacterial antigens. Electron microscopy and genome sequencing were performed to classify the isolated phage, and the host range was determined by spot tests and plaque assays. One-step growth curves and time-kill assays were conducted to characterize the life cycle of the phage, and to interrogate the multiplicity of infection (MOI) necessary for killing. A mouse model of infection was used to determine whether the phage could be used therapeutically against EAEC *in vivo*. Anaerobic culture in the presence of human fecal bacteria determined whether the phage could kill EAEC *in vitro*, and to assess whether the microbiome had been altered.

**Results:** The isolated phage, termed *Escherichia virus PDX*, is a member of the strictly lytic *Myoviridae* family of viruses. Phage *PDX* killed EAEC isolate EN1E-0227, a case-associated isolate from a child in rural Tennessee, in a dose-dependent manner, and also formed plaques on case-associated clinical EAEC isolates from Columbian children suffering from diarrhea. A single dose of *PDX*, at a MOI of 100, one day post infection, reduced the population of recovered EAEC isolate EN1E-0227 bacteria in fecal pellets in a mouse model of colonization, over a five-day period. Phage *PDX* also killed EAEC EN1E-0227 when cultured anaerobically *in vitro* in the presence of human fecal bacteria. While the addition of EAEC EN1E-0227 reduced the α-diversity of the human microbiota, that of the cultures with either feces alone, feces with EAEC and *PDX*, or with just the *PDX* phage were not different statistically, as measured by Chao1 and Shannon diversity indices. Additionally, β-diversity and differential abundance analyses show that conditions with *PDX* added were not different from feces alone, but all groups were significantly different from feces + EAEC.

**Conclusions:** The strictly bacteriolytic, *Myoviridae Escherichia virus PDX* killed EAEC isolate EN1E-0227 bacteria both *in vivo* and *in vitro*, while simultaneously not altering the diversity of normal human microbiota in anaerobic culture. Thus, the *PDX* phage could be part of an effective therapeutic intervention for children in developing countries who suffer from acute, or persistent EAEC-mediated diarrhea, and to help reduce the serious effects of environmental enteropathy. Because the emerging pathogen EAEC is now the second leading cause of traveler’s diarrhea, *PDX* could also provide therapeutic relief for these individuals, particularly in light of the growing crisis of antibiotic resistances.

## Introduction

Diarrheagenic *Escherichia coli* are a major cause of infectious disease worldwide, and in particular for those living in developing countries due to decreased access to safe water and healthcare. Of the *E. coli* pathotypes, enteroaggregative *E. coli* (EAEC) is a heterogeneous category, causing both acute and persistent diarrhea (1). EAEC has been implicated in endemic and epidemic outbreaks of disease (2, 3), and is also the second leading cause of traveler’s diarrhea. This category of pathogen has been linked to persistent diarrhea and malnutrition of children and HIV/AIDS patients living in developing countries. In the US, it was the most commonly isolated diarrheal pathogen in emergency rooms in a large-scale study in Maryland and Connecticut (3). Thus, EAEC is an emerging pathogen, for those living both in the developing and more developed regions of the world.

Antibiotics are generally prescribed for traveler’s diarrhea, but with the ever-increasing problem of drug resistance, treatment options are becoming more limited (4). The type of antibiotic given will depend on regional resistance patterns (5). In the developing countries, in addition to causing life-threatening fluid loss in children, recurrent and persistent bouts of diarrhea caused by these bacteria can result in nutrient and growth deficiencies (6). Though ultimately self-limiting, due to the persistent nature of EAEC disease, the syndrome is not effectively controlled by oral rehydration therapies alone. Thus, combined with the problem of drug resistance (7), alternate treatment strategies are urgently needed.

Microbiologists are generally familiar with the work of Felix d’Herelle, who discovered bacteriophage during an outbreak of dysentery among soldiers during World War I, and the subsequent success of phage therapy against bacterial infections in eastern Europe in the subsequent decades (8, 9). However, today even those in the general public are learning of phage therapy due to the prevalence of multiple drug resistant (MDR) bacteria. More recently, phage have been used successfully for treatment of many bacterial pathogens, for example, in Poland (10). However, few phage therapy clinical trials have been conducted in Western countries. Two successful phage therapy applications, of single dose phage cocktails, a mixture of bacteriophage that target different pathogenic bacteria, have been conducted in the US. In one case, the phage cocktail was administered topically to burn victims in a military hospital in Queen Astrid military hospital in Brussels, Belgium (11). The other successful Phase II clinical trial of phage therapy was administered to patients suffering from chronic MDR *Pseudomonas aeruginosa* ear infections (12). This study was terminated early as the treatment group exhibited such impressive recoveries that the physicians wanted to be able to deliver the phage preparation to the control group of patients to treat their infections. Three patients enrolled in the study experienced full resolution of symptoms as determined by both the study physicians and self-reporting. Though there are significant hurdles for developing mechanisms for FDA approval of the clinical use of bacteriophage (13, 14), the results of these studies strongly suggest that the efficacy of phage therapeutics is not the limiting factor in making such applications available.

A clear advantage of using phage as therapy against bacterial infections is that they are thought to only target specific pathogens, often only one species or strain, without disrupting the normal microbiota, or causing dysbiosis. This is in stark contrast to more traditional antibiotics, which are life-saving drugs, but are also guaranteed to cause dysbiosis due to the conserved nature of drug targets. Recent reports demonstrate that this type of disruption can be associated with a number of different deleterious health effects (15). One such review presents data on the loss of microbiome diversity in children treated with antibiotics (16), which includes a reduction of α-diversity, a metric of biodiversity, of up to 40% from ciprofloxacin, an antibiotic commonly given for EAEC infections. While there have been several studies to demonstrate intestinal microbiome dysbiosis due to antibiotic treatment in human volunteers (17), only a few have shown that phage can be used to treat bacterial infections without causing dysbiosis (18–20).

In this study, using a sequential isolation technique to enrich for bacteriophages capable of infecting the desired hosts, we isolated a strictly lytic phage from wastewater in Portland, OR, that kills strains of EAEC bacteria. By electron microscopy techniques, the *PDX* phage was tentatively identified as a member of the *Myoviridae* family that infect *E. coli*, and this was confirmed through functional genome bioinformatic and phylogenetic analysis. We characterized the growth and infectivity of the *PDX* phage, and showed that it kills a case-associated EAEC strain both *in vitro* and *in vivo*. In a mouse model, a single dose of the phage one day post infection significantly reduced the number of bacteria recovered from feces over a five-day period. Lastly, we show that *PDX* kills EAEC when cultured anaerobically in the presence of human fecal bacteria, without altering the α-diversity of the human microbiota.

## Materials and Methods

### Bacterial strains and growth conditions

Bacteria used for phage isolation and clinical EAEC isolates for host susceptibility testing are listed in Table 1. EPEC clinical isolates for host susceptibility testing are listed in Table S1. All strains were grown in Lysogeny Broth (LB) at 37°C with shaking at 225 RPM or on LB agar at 37°C. All phage lysates were stored in SM gel buffer (50 mM Tris-HCl [pH 7.5], 0.1 M NaCl, 8 mM MgSO_4_, 0.01% gelatin) at 4°C, and warmed to room temperature for all experiments unless otherwise stated.

**Table 1.** Escherichia coli strains used in this study.

| Strain | Genotype/<br>Serotype | Syndrome<sup>1</sup> | Spot Test<br>Result<sup>2</sup> | Plaque<br>Formation<sup>3</sup> | Source |
| --- | --- | --- | --- | --- | --- |
| EN1E-0227 | EAEC<br>O86:H27 | D | + | + | (65) |
| LRT9 | EPEC<br>O11:abH2 | U | + | + | (47) |
| MC4100 | *F- araD139*<br>*Δ(argF- lac)*<br>*U169 rpsL*<br>*150 StrR*<br>*relA1*<br>*flhD5301*<br>*deoC1 ptsF25*<br>*rbsR* | NP | + | + | (66) |
| BC/Ae017.2 | EAEC<br>AM<sup>R</sup>, AMC<sup>R</sup> | D | - | - | UB<sup>4</sup> |
| BC/AaUIM099.2 | EAEC | D | + | - | UB |
| BC/AaUIM171.3 | EAEC<br>AM<sup>R</sup> | D | - | - | UB |
| BC/AaUIM206.1 | EAEC | D | + | - | UB |
| BC/Ac277.1 | EAEC | D | - | - | UB |
| BC/Ac338.3 | EAEC<br>AM<sup>R</sup>, AMC<sup>R</sup> | D | + | + | UB |
| BC/AaUIM363.2 | EAEC | D | + | + | UB |
| BC/Ac408.1 | EAEC | D | + | - | UB |
| BC/Ac422.2 | EAEC | D | - | - | UB |
| BC/Ac482.1 | EAEC<br>AM<sup>R</sup>, SXT<sup>R</sup> | D | - | - | UB |
| BC/AaUIM517.1 | EAEC | D | - | - | UB |
| BC/Ac512.2 | EAEC | D | - | + | UB |

<sup>1</sup>D, acute diarrhea; U, details unknown; NP, non-pathogenic lab strain.<sup>2</sup>Plus sign indicates strain is killed by *Escherichia virus PDX*.
<sup>3</sup>Plus sign indicates *PDX* replicates (can form plaques on) the strain.
<sup>4</sup> Division of Pediatrics Infectious Diseases, University at Buffalo.

### Phage isolation and propagation

Bacteriophage *PDX* was initially isolated from untreated (influent) sewage collected at the Portland Wastewater Treatment Plant on October 1, 2015. The raw sewage was filtered through a 0.2 µm filter (Corning, Corning NY) to remove bacteria. The filtered sewage was then added to host bacteria in liquid culture. Phage-host solutions were combined with 0.5% LB soft agar and overlaid on LB agar plates. Plates were then observed for the presence of plaques after overnight growth at 37°C. Clear, defined plaques of interest were isolated and transferred using a sterilized inoculating needle and agitation in 1 mL solution of lambda diluent (10mM Tris, pH 7.5; 10 mM MgSO_4_). To obtain a high titer phage preparation, a crude lysate was generated. A clear *PDX* plaque was isolated and suspended in 1 mL of sterile lambda diluent. The phage suspension was combined with overnight culture of the bacterial host and incubated at 37°C with shaking at 225 RPM overnight. Chloroform was added to kill any remaining bacteria. This solution was centrifuged at 4°C at 4000 x g for 10 minutes to remove bacteria. The supernatant was recovered and treated again with chloroform.

Phage were sequentially isolated from *E.coli* strain MC4100, EPEC strain LRT9, and EAEC clinical isolate EN1E-0227 using a sequential isolation technique to enrich for core antigen recognition (21). Several clearly defined plaques from the final overlay were picked using a sterile inoculating needle and suspended in microcentrifuge tubes containing SM buffer. The phage suspension was incubated at room temperature for 15 minutes to allow the phage to diffuse through the medium, then filtered through a sterile 0.2 µm filter to remove any bacteria that may have been transferred from the overlay plate. A host susceptibility assay was performed as described below to test the ability of each selected plaque to lyse strains MC4100, LRT9, and EAEC EN1E-0227. The final phage suspension was then propagated on EAEC strain EN1E-0227 to make a purified high titer lysate. Bacteria were removed from the culture by centrifugation at 6,055 x g, then the supernatant was sterile filtered with a 0.2 µm filter. The titer of the lysate was determined by serial dilutions and overlay assay as described. After overnight incubation, the plates were observed and the number of plaque forming units per mL (PFU/mL) was calculated. The purified high titer lysate in SM buffer was stored at 4°C.

### Host susceptibility assay

A spot test was used to determine the host range of *PDX* on clinical EAEC and EPEC clinical isolates (Table 1; Table S1). Each LB agar plate was streaked with 3-4 spatially separated bacterial strains using a sterilized inoculating loop. A 10 μl purified lysate sample was pipetted on top of the center of each linear bacterial streak and incubated overnight at 37°C. Host susceptibility was indicated by a clearing within the streak of a given bacterial strain. Clinical EAEC and EPEC isolates were also tested for the ability to support *PDX* replication using the overlay assay described above and observing phage plaques (Tables 1 and S1).

### Electron microscopy

*PDX* was analyzed using transmission electron microscopy (TEM). Purified high titer lysate was precipitated in 25% PEG 6000-8000 in 2.5M NaCl and stored overnight at 4°C. Precipitated phage solutions were centrifuged at 4°C, 4000 x g for 10 minutes. The supernatant was removed and the phage pellet was suspended in 40µL of 50 mM Tris, pH 7.4, 10 mM MgSO_4_. Phage suspension was placed on copper coated formvar grids (Ted Pella, Inc., Redding, CA) and negatively stained with 1% uranyl acetate. Samples were examined under a FEI Tecnai™ Spirit TEM system at an operating voltage of 120 kV (FEI, Hillsboro, OR). All transmission electron microscopy was performed at the Multi-Scale Microscopy Core (MMC) with technical support from the Oregon Health & Science University (OHSU)-FEI Living Lab and the OHSU Center for Spatial Systems Biomedicine (OCSSB).

### Phage genome sequencing and bioinformatics

A high titer lysate was purified using a cesium chloride gradient, then genomic dsDNA was extracted with an Epicentre DNA (Madison, WI) isolation kit according to the manufacturer’s instructions. All library preparation and Illumina next generation sequencing was performed at the Massachusetts General Hospital (MGH) Genome Core (Cambridge, MA). Raw reads were filtered and processed for quality using Galaxy (22). The read adaptors were trimmed with Trimmomatic Galaxy (v. 0.32.3) using the default settings (23). The quality of the trimmed reads was processed with Fast QC (v. 0.11.4) using default settings (24). The reads were assembled with a de novo De Bruijn algorithm-based software SPAdes with the virus kingdom selected (v 3.7.1) (25). Next the quality of the assembly was determined using the Quality Assessment Tool for genome assemblies (QUAST v 2.3) (https://www.bioinformatics.babraham.ac.uk/projects/fastqc/). The assembled genome was annotated using ORF annotation (NCBI ORFfinder) and PROKKA with default settings and National Center for Biotechnology Information (NCBI) smart Basic Local Alignment Tool (BLAST) (v 1.12) (26). Annotation outputs were compared in tandem and putative coding sequences (CDS) and proteins with known functions above 95% were selected using the default algorithm parameters.

Next, output CDS sequences from the genomic annotations were aligned for similarity to other sequences in the National Center for Biotechnology Information (NCBI) database using the Basic Local Alignment Search Tool (BLAST; v. 2.3.1) suite, specifically using the blastp and tblastn tools with default settings (27, 28). Finally, genes associated with structural components were analyzed using the Virfam webserver, which uses a machine learning-hierarchical agglomerative clustering method to construct a protein phylogeny between *PDX* and all other phages contained in the ACLAME database.

### Accession number

The assembled genome sequence of the *Escherichia* virus *PDX* was deposited in GenBank under accession number MG963916.

### One-step growth curves

As our target for therapy is EAEC, growth kinetics of *PDX* using the one-step growth curve method were performed on strain EN1E-0227. LB liquid cultures were grown to an OD_600_ of 0.6-0.7, mid to late exponential phase. The purified phage lysate was added to the liquid culture tubes to infect the bacteria with a multiplicity of infection (MOI) of 0.01. The liquid bacterial culture and phage lysate were incubated at room temperature for 10 minutes. Following the incubation period, the solution was centrifuged at 8,000 x g for 10 minutes. The supernatant was discarded and the pellet was re-suspended in LB to make the absorption mixture. Immediately after re-suspension the absorption mixture was serially diluted and plated with the bacterial host. The phage was incubated within a standard plaque-overlay. This procedure was repeated at 20-minute time intervals for a total of 120 minutes. After 24 hours of incubation, the plaques on all the plates were recorded and the PFU/mL was calculated. The titer over time (120 minutes) was plotted to obtain a one-step growth curve.

### Time-Kill Assays

Bacterial density was measured over time to determine the course of phage infection. Liquid cultures of EAEC strain EN1E-0227 were incubated with *PDX*. The *in vitro* lytic efficiency of the phage was examined at several MOIs. Wells of a flat-bottomed 96-well microtiter plate were filled with 100 μl of inoculated double-strength LB and 100 μl of purified lysate dilutions in SM buffer. Bacterial cultures were grown to late exponential phase and diluted to 10^6^ CFU/mL. *PDX* lysate was diluted to 10^5^, 10^6^, 10^7^ or 10^8^ PFU/mL, corresponding to MOIs of 0.1, 1, 10, and 100, respectively. Each phage-host combination at specific MOIs was performed in triplicate wells, and each experiment was performed three times. Controls for plate sterility, phage suspension sterility, and bacterial growth without phage addition were included. To prevent between-well contamination by aerosolizing of bacteria, an adhesive transparent plate cover was placed over the top of the plate. The plates were incubated at 37°C with orbital shaking for 12 hours and OD at 600 nm was measured using a Microtiter Plate Reader (Sunrise, Tecan Group Ltd., Austria) at 30-minute intervals.

### Murine model of intestinal colonization

C56BL/6 mouse models have been previously developed for EPEC (29) and EAEC (30, 31). Although mice do not display robust histopathological or clinical signs of disease, they serve as an *in vivo* colonization model to extend findings from *in vitro* experiments. C57BL/6J WT mice aged 6 – 8 weeks received ampicillin (1 g/L) added to the drinking water for two days to reduce the gut microbiome (30). After two days, ampicillin water was removed and replaced with sterile drinking water for the rest of the experiment. Mice were infected via oral gavage with 4.0x 10^7^ CFU/mL *E. coli* resuspended in 100 µL of 1X PBS, pH 7.4 for an effective dose of 4.0x 10^6^ CFU/mouse (29). Fifteen minutes prior to inoculation with bacteriophage *PDX*, mice were given 100 µL 2.6 % sodium bicarbonate by oral gavage to neutralize stomach acid (17). Bacteriophage were diluted to 4.0x 10^9^ PFU/mL in SM buffer + 0.01% gelatin and 100 µL were introduced by oral gavage for an effective dose of 4.0x 10^8^ PFU/mouse. EAEC strain EN1E-0227 was monitored by daily collection of feces and enumeration of CFU/g feces. Feces were resuspended and serially diluted in LB plus 1% saponin to reduce bacterial clumping and plated on LB plates containing 100 µg/mL streptomycin. Streptomycin-resistant bacteria from the mice tested indole positive, demonstrating EAEC was successfully isolated, cultured and enumerated.

### Ethics Statement

C57BL/6J WT mice (male and female) were purchased from Jackson Laboratories and housed at Oregon Health and Science University (OHSU) under specific pathogen free conditions in an ABSL-2 facility in accordance with University and Federal guidelines. All experimental protocols were approved by OHSU’s Department of Comparative Medicine (DCM) in accordance with the University’s Institutional Animal Care and Use Committee (IACUC).

### Anaerobic culture

Fecal cultures were prepared using methods adapted from Caldwell and Bryant’s 1966 Hungate roll tube method for non-selective growth of rumen bacteria (32). In order to mimic the nutrient, mineral, pH, and environmental conditions of human gut *in vitro*, M10 broth media was prepared and dispensed into sterile Hungate tubes with rubber stoppers. Anaerobic conditions were achieved by sparging with 100% CO_2_ gas at 10 PSI for 15 minutes. Resazurin dye was used as a visual indicator of anaerobic conditions. In the presence of oxygen, resazurin is reduced to resorufin, a distinctly red compound. In absence of oxygen (*i.e*. anaerobic conditions) resazurin turns M10 media colorless (32).

Feces were collected from a 21-year-old male in Portland, Oregon who had not received antibiotics for over one year. One gram of fresh feces was immediately placed in a sterile Hungate tube and bubbled for 10 minutes at 10 PSI with 100% CO_2_ gas. The feces were suspended in 10 ml of sterile M10 medium, which was transferred aseptically and anaerobically via a sterile syringe and hypodermic needle from one stoppered Hungate tube to another. The resulting slurry was vortexed vigorously and serially diluted in stoppered Hungate tubes of M10 to a final concentration of 1.0×10^-4^ g feces /mL, and then grown anaerobically for 16 hours at 37° C using the Hungate method (32). After a 16-hour incubation period, the starting culture was used to inoculate subsequent fecal cultures as described below.

Streptomycin-resistant (strR) EAEC EN1E-0227 were grown anaerobically under 100% CO_2_ with fecal slurry and *PDX* using the Hungate method in M10 medium (32). Cultures were inoculated to final concentrations: 16-hour fecal, M10 culture diluted 1:100, 4.0 x 10^3^ CFU/mL EAEC of isolate EN1E-0227, and 4.0 x 10^5^ PFU/mL phage for a MOI of 100. Cultures were incubated for 16 hours in a stationary incubator at 37°C. DNA from these cultures was isolated for 16S rDNA sequencing analysis after 16 hours of incubation.

Immediately prior to DNA extraction, a sample from each culture was plated to enumerate the concentrations of strR EAEC in the anaerobic cultures. As a control, fecal samples were plated on LB agar with streptomycin, and no growth was observed. Serial dilutions were performed in a 96-well plate in sterile LB, then plated in triplicate, 10-µL drops, of each concentration on LB-agar containing 100 µg/mL streptomycin. Plates were incubated for 20 hours at room temperature and colonies were counted using a dissection microscope.

DNA was isolated from fecal cultures using a GenElute Bacterial Genomic DNA Kit (Sigma-Aldrich, St. Louis, MO) following the protocol for cultures containing gram-positive and gram-negative bacteria. Whole genomic DNA was quantified using a Nanodrop 1000 (Thermo Fisher Scientific, Waltham, MA).

### 16S Metagenomic sequencing and analysis

Genomic DNA was sent to the Oregon State University Center for Genome Research and Biocomputing for multiplex sequencing analysis using the Illumina MiSeq sequencing platform, with paired-end forward and reverse reads of 150 bp in length and a sequencing depth of coverage totaling 1 million reads. The V3-V4 region of the 16S rRNA gene was amplified and used to prepare a dual index library using the Nextera XT library preparation kit. Sequences were quality filtered and demultiplexed using FastQC, then processed in RStudio (version 1.1.383, 2015, RStudio: Integrated Development for R. RStudio, Inc., Boston, MA URL http://www.rstudio.com/). using the DADA2 package (33)to de-noise and merge the forward and reverse reads. As per the DADA2 pipeline, chimeric sequences were removed and pre-processed reads were clustered into 100% identical sequences representing amplicon sequence variants (ASVs) to capture the full extent of genetic diversity in the microbiome (34), then read counts were converted into a count table. ASVs were assigned taxonomy to the level of genus using the SILVA ribosomal RNA gene database (release 132) as a reference (33, 35). Non-bacterial taxa were removed from the dataset. ASV sequences were aligned using ClustalOmega (version 1.2.4) (36)and a 16S phylogenetic tree was constructed using FastTree (version 2.1.7) (37).

The count table, taxonomy table, metadata table, and phylogenetic tree were used for quantitative and qualitative analysis using the R-packages Phyloseq (version 1.24.2) (38) and DESeq2 (version 1.20.0 (39)). Chao1 and Shannon diversity indices were calculated in Phyloseq from unnormalized ASV counts as measures of alpha diversity, using Tukey Post-Hoc HSD for statistical analysis. Chao1 is an estimated measure of species richness that adjusts for variable read depths across samples using the number of singletons in each sample (40, 41), while the Shannon index is an estimate of diversity accounting for both richness and evenness (42). ASV counts were normalized across samples using a variance-stabilizing transformation (VST) implemented in DESeq2 prior to calculating beta diversity. The adjusted VST count table and phylogenetic tree were used to construct a weighted-UniFrac (43) distance matrix, and beta diversity was plotted using principal coordinate analysis (PCoA) as the ordination method. To test whether samples form significantly different clusters based on treatment groups, the Adonis2 tool from the Vegan package (version 2.5-2) (44)was used to run a PERMANOVA (45) for testing whether group centroids were significantly different. Finally, to test which taxa are differentially abundant across treatment groups, pairwise binomial Wald tests were performed on VST adjusted data in DESeq2. Each statistical test described above was executed a second time on the dataset with ASVs representing the *Escherichia* genus filtered out to confirm any microbiome community differences detected were not primarily due to counts of inoculated bacteria.

## Results

### Isolation and initial characterization of a *Myoviridae* phage

Phages were isolated from clear, ∼2 mm plaques on LB agar that formed against *E. coli* host strains when overlaid with filtered, raw sewage. *Escherichia virus PDX* was initially isolated against *E. coli* strain MC4100 because the altered LPS in laboratory strains allows for phage recognition of more conserved, core antigens (46). Similarly, EPEC LRT9 was used because it is also a laboratory strain, amenable to P1 transduction (47). The phage stock prepared from these plaques was purified using a sequential host isolation technique adapted from Yu *et al*., 2016 (21). Sequential host isolation with MC4100, then EPEC and EAEC strains LRT9 and EN1E-0227, respectively, allowed us to enrich the phage for core antigen recognition. The lysates for *Escherichia virus PDX* had a titer of 8.92×10^12^ PFU/mL when propagated on EAEC strain EN1E-0227, the host bacterial pathogen for this study. As a negative control, the phage was unable to lyse a strain of *Bacillus subtilis* (ATCC #6051).

To test how broadly *PDX* recognized EAEC bacteria we found that the phage lysed five and formed plaques on two of 12 EAEC case-associated strains from children in Columbia (Table 1). One of these strains, BC/Ac338.3 exhibited multiple drug resistances, to AM (ampicillin) and AMC (amoxicillin clavulanate). *PDX* lysed nine and form plaques on five out of 20 clinical, diarrheal EPEC isolates from children living in the Seattle area (Table S1; (48)). These enteropathogenic isolates represented a range of serotypes and variety of syndromes, including acute, chronic, and in one case, strain TB96A, serotype O75:HN, bloody diarrhea. Thus, the *Escherichia virus PDX* replicated in, and formed plaques on multiple case-associated EAEC and EPEC isolates.

To obtain preliminary classification of the *PDX* phage we employed transmission electron microscopy (TEM). TEM imagining revealed the phage most likely belonged to the family *Myoviridae,* identified by their canonical morphology: a small-isometric non-enveloped head with a long contractile tail (Fig. 1; (49)). The head diameter of *PDX* was 76.4 ± 4.34 nm and the tail length was 114.0 ± 2.26 nm (n = 13) (Figs. 1ABC). Based on morphological features and metrics obtained using TEM, we tentatively identified *PDX* to be a member of the *Myoviridae* family that infect *E. coli* (Fig. 1D).

**Figure 1.**
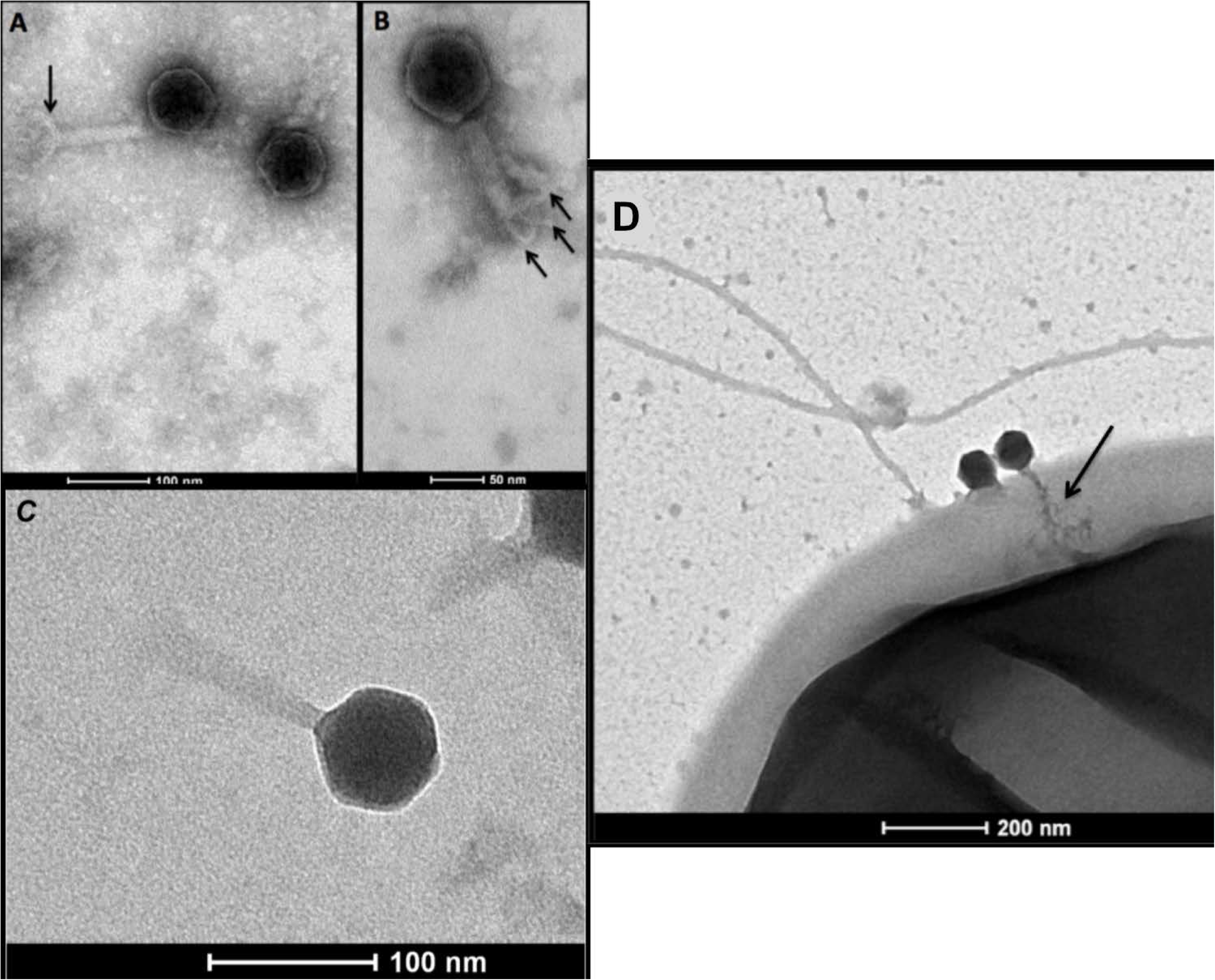
Transmission electron micrographs of bacteriophage *Escherichia virus PDX*. Samples were negatively stained with 1% uranyl acetate solution, prepared on a carbon formvar grid, and were imaged at 120 kV on a FEI Tecan-i Spirit TEM system. The arrow in (A) indicates the base plate. The arrows in (B) identify three putative tail fibers. The image in (C) illustrates the small isometric, non-enveloped head with long contractile tail, specific to the family *Myoviridae*. (D) illustrates *PDX* infecting EAEC, with contractile tail, indicated by arrow, penetrating the outer membrane and periplasmic space. Scale bars are as indicated.

### Genomic sequence analysis

Therapeutic phage must be virulent, or lytic, as to minimize risks associated with horizontal gene transfer by temperate, or lysogenic phage (50). Thus, we used genome sequence analysis to confirm that *PDX* was a member of the strictly lytic family of *Myoviridae* phages (51). The genome of *PDX* is 138,828 bp dsDNA, typical of the *Myoviridae* family of viruses. The GC content is 43%, which is comparable to other lytic phages (FastQC Galaxy). We identified 206 putative coding domain sequences (CDS) (PROKKA, ORFfinder), but did not identify toxin-encoding or lysogenic genes, which are characterized as any gene containing the terms “integrase”, “excisionase”, “recombinase”, or “repressor” within the genome when compared to the NCBI BLAST database. Additionally, in a 20-standard amino acid (aa) search, a total of 6 putative tRNAs were predicted with open reading frames (ORFs) ranging from 72-88 bp in length (tRNAscan-SE). Virfam software uses the translated amino acid (aa) of CDS from a phage genome as input and searches for similarity to other structural phage protein sequences deposited in the Aclame database.

Based on the results of the Virfam analysis, the head-neck-tail structure genome organization in *PDX* belongs to the Neck Type One – Cluster 7 category (Fig. 2A). Neck Type One phage genomes contain the following proteins: portal protein, Adaptor of type 1 (Ad1), Head-closure of type 1(Hc1), Neck protein of type 1(Ne1), and tail-completion of type 1 (Tc1) (Fig. 2B). Cluster 7 phage genomes are reported in Virfam to be strictly *Myoviridae* with relatively small genome sizes and gene content (61-150 genes), though *PDX* contains 206 CDS (Table 2).

**Figure 2.**
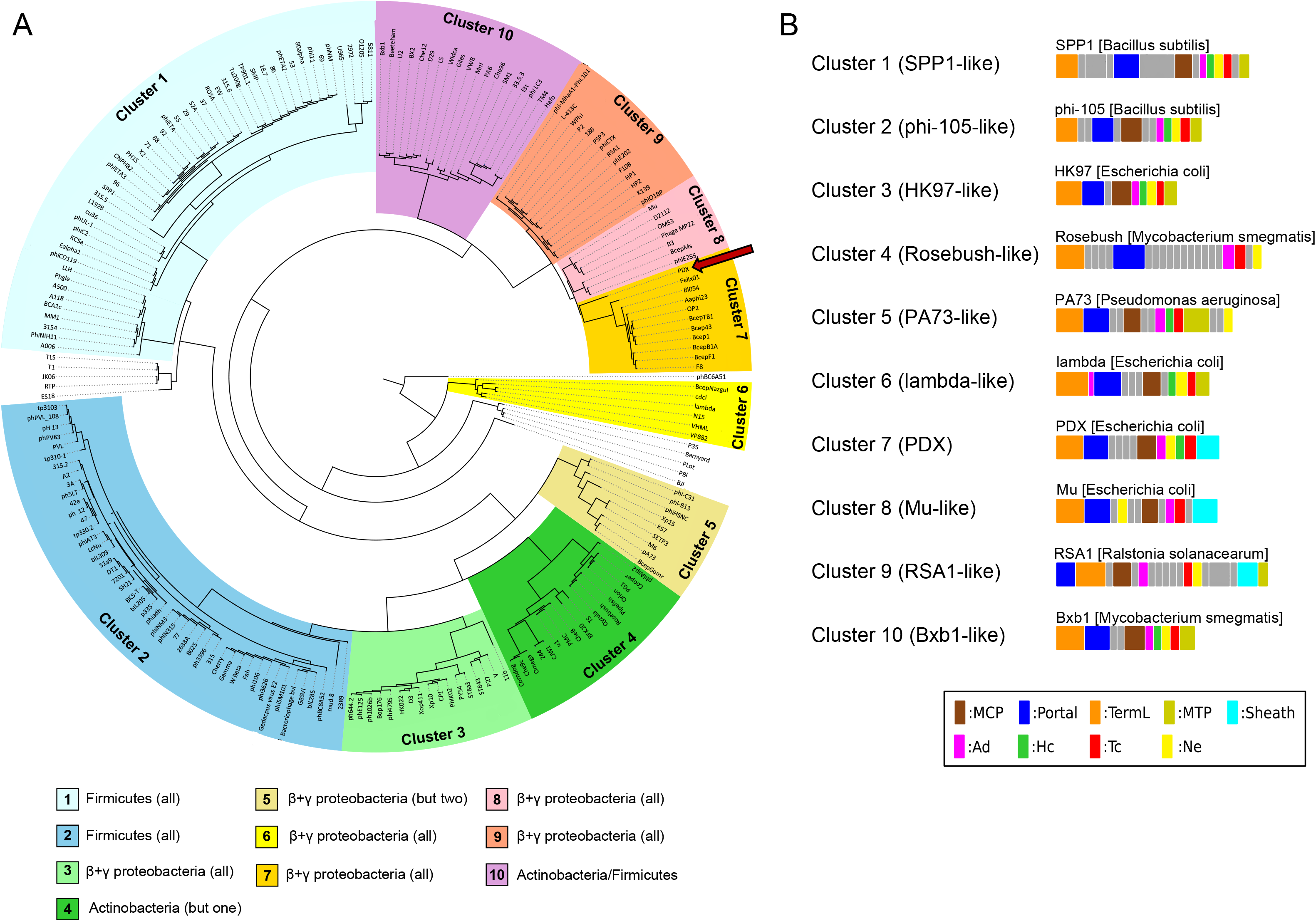
Genome and structural protein classification analysis of *PDX*. (A) Tree representation of *PDX* (indicated by red arrow) classification among other Caudovirales phages, constructed from a hierarchical agglomerative clustering procedure applied to a matrix of similarity scores between pairs of phages (by combining HHsearch probabilities with the percentage identity) and present in the Aclame database. The different branches of the tree were sorted into 10 Clusters and are highlighted by different background colors. The bacterial host(s) of the phages are listed for each Cluster in the legend at the bottom of the figure. (B) Gene organization of *PDX* and other phage genomes that belong to the Neck One Clusters. The legend at the bottom indicates to which family a protein belongs. In the legend, “but one” and “but two” indicate that all but one or two of the phages are lytic against the bacterial family indicated by the colored boxes below the phylogeny.

**Table 2.** Genomic analysis of *PDX*

| Feature | Quantity |
| --- | --- |
| Size (bp) | 138,828 |
| GC content (%) | 43 |
| Coding domain sequences (CDS) | 206 |
| tRNAs | 6 |

By BLAST analysis, the three genomes with the highest whole-genome nucleotide identity match were *Escherichia coli* O157 typing phage 4 (98%), *Escherichia* Phage JES-2013 (98%), and *Escherichia* Phage Murica (Table S2). Murica is a *Myoviridae* phage that shares a 97% nucleotide identity with *PDX*, and is strictly lytic against enterotoxigenic *E.coli* (ETEC) (52). *E. coli* O157 typing phage 4 is lytic against *E. coli* O157, and also shares 99% amino acid identity with the *PDX* tail fiber proteins (53). Phage JES-2013 is lytic against *Lactococcus lactis* (54). The putative tail fiber sequences from *PDX* had a 99% amino acid identity with *Myoviridae Escherichia* phage V18 (55). Overall, the phage that had the highest percent identity scores with the *PDX* genome sequence and tail fiber amino acid sequence were all strictly lytic *Myoviridae* against various *E. coli* strains.

### Phage *PDX* kills enteroaggregative *E. coli* both *in vitro* and *in vivo*

To characterize the phage life cycle, we performed a one-step growth curve of *PDX* (Fig. 3A). *PDX* infecting EAEC strain EN1E-0227 had a maximum relative virus count of 3.60×10^6^ PFU/mL after 100 minutes of infection. The bacteriolytic activity of *PDX* against EAEC strain EN1E-0007 was measured in a time-kill assay revealing a dose-dependent inhibition of bacterial growth. At a MOI of 100, *PDX* severely suppressed growth of EAEC (Fig. 3B). Bacteria that were not treated with phage showed normal logistic growth curves that match the expected growth rate for this strain. A dose dependent relationship between MOI and point of infection was observed: at greater MOI, infection occurred earlier than for lower MOI (Fig. 3B).

**Figure 3.**
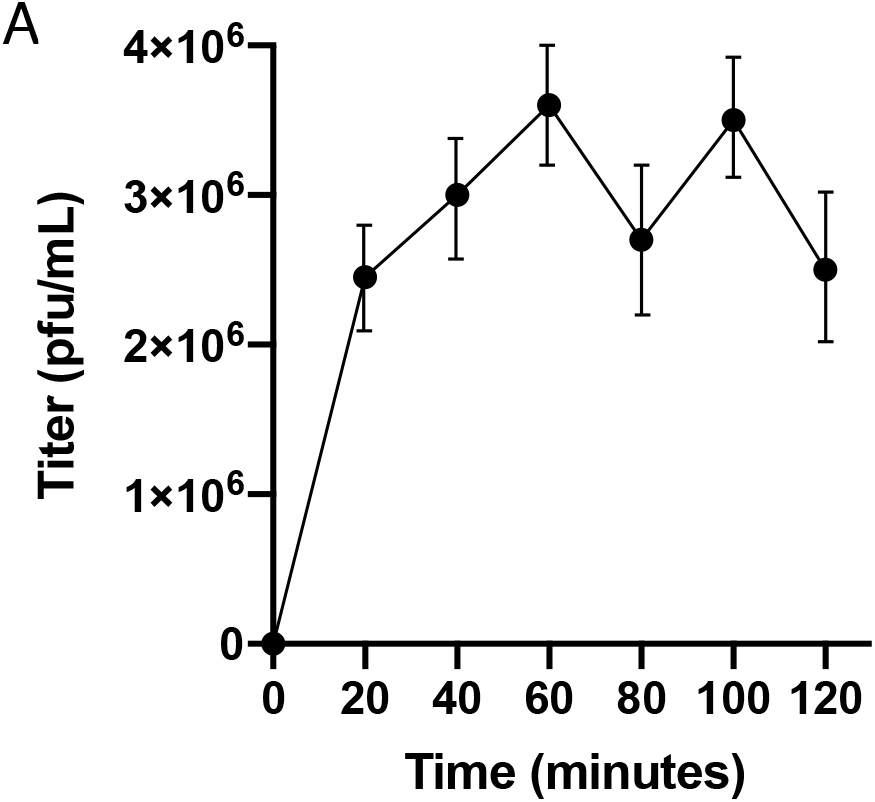
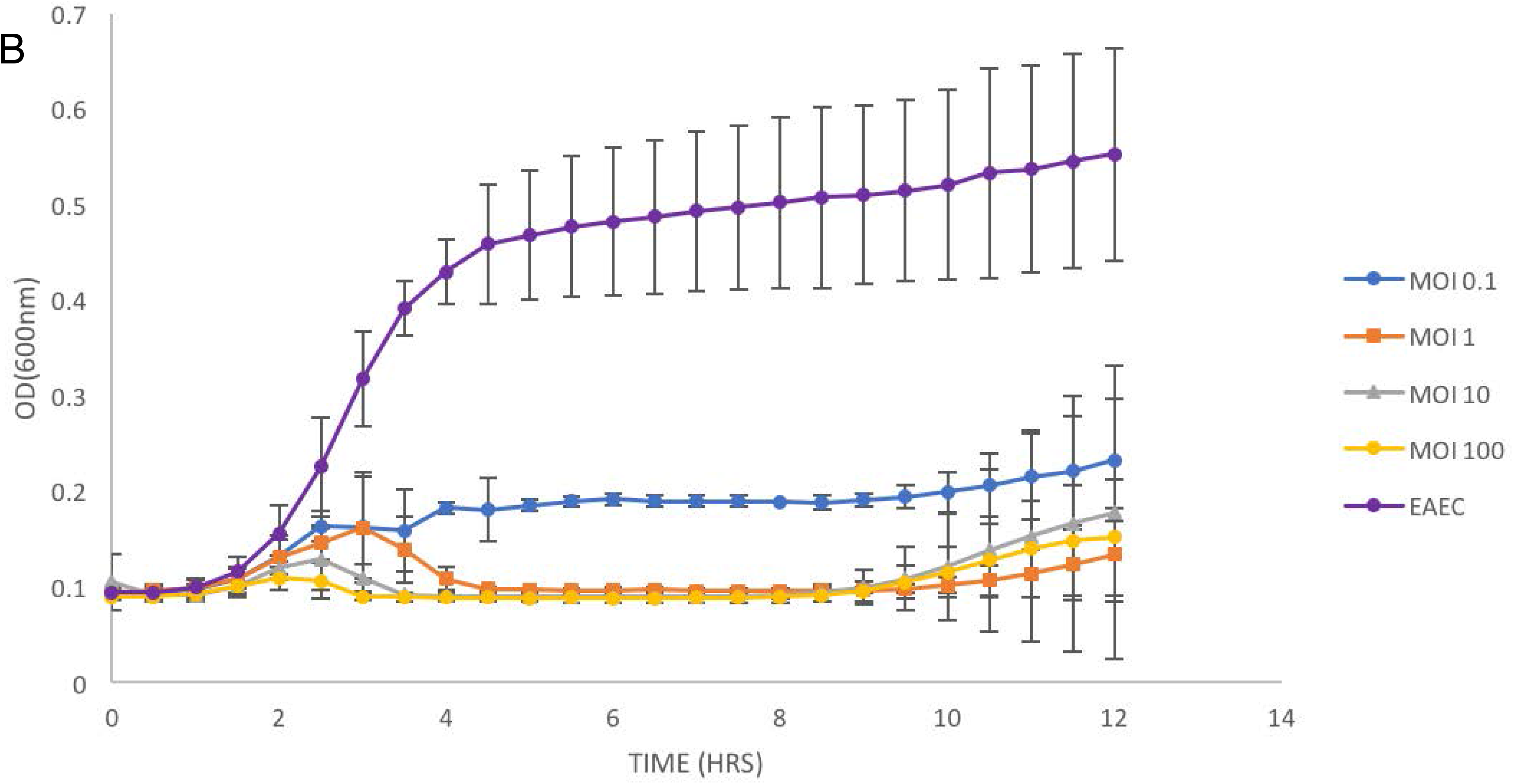
One-step growth curve analysis and time-kill assays of EAEC infected with *PDX.* (A) Over the 120-minute infection period, bacterial cultures were sampled, serially diluted and plated with a plaque-overlay procedure (see Materials and Methods). After 24-hour incubation, the plaques were counted and the titer (PFU/mL) was determined. Error bars represent the standard error of the mean. (B) Time-kill assay of EAEC strain EN1E-0227 treated with *PDX*. In a 96-well plate, exponential culture was inoculated with dilutions of *PDX* high titer lysate for MOI (multiplicity of infection) ranging from 0.1, 1, 10 and 100. The plates were incubated at 37°C with orbital shaking for 12 hours and OD of 600 nm was measured using a Microtiter Plate Reader (Tecan). Two independent assays were performed in triplicate, with representative, averaged assay data presented. Error bars indicate ±1 standard deviation.

In many phage treatments, an increase in bacterial growth was observed around 8-10 hours of incubation. This is likely due to the emergence of mutant bacteria that are resistant to phage infection and has been observed in other, similar studies (31, 56). We isolated the bacteria resistant to phage infection. Phage induction, by treatment with Mitomycin C, did not induce *PDX* plaques in the resistant, mutant EAEC. This result also supported the conclusion that *PDX* is strictly lytic, and appropriate for use as a therapeutic agent.

We next used a mouse colonization model to test the prediction that *PDX* can kill EAEC strain EN1E-0227 *in vivo*. Mice were given ampicillin in their drinking water one day prior to infection in order to compromise colonization resistance and help to establish infection by the pathogen. A single dose of 4 x 10^6^ CFU EAEC in PBS was introduced to all EAEC-challenged mice by oral gavage after antibiotics were removed from the drinking water. One day post infection, fecal pellets were collected and a single dose of 4 x 10^8^ PFU *PDX* was administered to all phage-challenged mice by oral gavage at a MOI of 100. CFU/g feces were enumerated across all subsequent days for the duration of the experiment. A single dose of *PDX* caused a statistically significant decrease in EAEC colony counts at two, three, and five days post infection (Fig 4). The most pronounced decrease in EAEC occurred at five days post in infection, with mean 8.9 log CFU/g feces in EAEC-challenged mice and mean 7.7 log CFU/g feces EAEC treated with *PDX* (*P* = 0.0016). As controls, we showed that the bacteria recovered from mouse feces were streptomycin resistant *E. coli*, by indole test, and that no bacteria from fecal pellets from uninfected mice grew on selective plates. We concluded that *PDX* reduced colonization of EAEC strain EN1E-0227 in the mouse model of infection.

**Figure 4.**
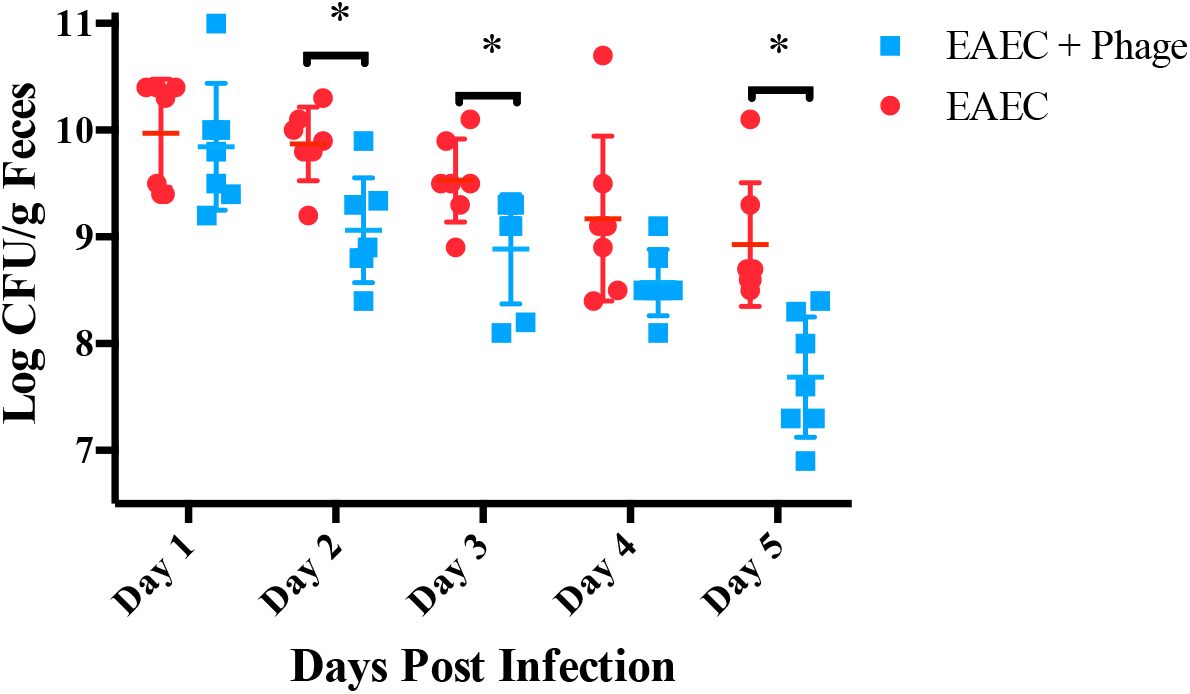
A single dose of *PDX* decreased murine gut carriage of EAEC. 24 hours after inoculation with EAEC, mice were gavaged with a MOI of 100 PDX phage or an equal volume of buffer control. No colonies were detected in phage-only control mice across the five-day period. Statistical analysis was carried out to compare *PDX*-treated mice to control treated mice at each indicated day (n =14 animals per day, Day 2, P=0.0097; Day 3, P=0.0214; Day 5, P=0.0016). Each error bar is constructed using 1 standard deviation from the mean. Levels connected by an asterisk are statistically significantly different (P<0.05).

### *PDX* phage kills EAEC strain EN1E-0227 without altering diversity of human fecal bacteria when propagated anaerobically *in vitro*

To determine whether *PDX* could kill EAEC bacteria in the presence of the human gut microbiome, feces were propagated anaerobically in M10 medium in the presence of the diarrheagenic EAEC strain EN1E-0227. Cultures were inoculated with 4.0 x 10^3^ CFU/ml EAEC or bacteria plus *PDX* phage 4.0 x 10^5^ PFU/ml, to a MOI of 100, with or without 25 μg/mL ciprofloxacin, an antibiotic commonly given to treat traveler’s diarrhea (57). Cultures lacking a *PDX* challenge grew from the 4.0 x 10^3^ CFU/ml inoculum to 9.0 x 10^7^ CFU/mL of EAEC in the absent of any additional agent (Table 3). In the cultures receiving EAEC and *PDX* phage the inoculum remained relatively unchanged, decreasing slightly to 1.7 x 10^3^ CFU/mL from the initial 4.0 x 10^3^ CFU/mL. The cultures with ciprofloxacin added to 25 μg/ml were killed, with no bacterial growth during enumeration (Table 3). We concluded that *PDX* phage reduced growth of EAEC strain EN1E-0227 by nearly 5 log CFU/ml in the presence of human microbiota when cultured anaerobically to mimic the human gut.

**Table 3.** Colony counts of streptomycin resistant EAEC strain EN1E-0227 in fecal culture after 16-hour incubation at 37 °C.

| Treatment | Feces (CFU/mL) | Feces + EAEC<sup>c</sup><br>(CFU/mL) |
| --- | --- | --- |
| Neat | 0 | $9.0 \times 10^7$ |
| *PDX*<sup>a</sup> | 0 | $1.7 \times 10^3$ |
| Ciprofloxacin<sup>b</sup> | N/A | 0 |

Human fecal samples were collected and cultured anaerobically under the following regime: Feces only, Feces + EAEC, Feces + EAEC + Ciprofloxacin, Feces + EAEC + *PDX* and Feces + *PDX*. Each challenge was cultured in triplicate.
*a*: Cultures were inoculated to $4.0 \times 10^5$ PFU/ml *PDX* for a MOI of 100.
*b*: Cultures were inoculated to 25 µg/mL Ciprofloxacin
*c*: Cultures were inoculated to $4.0 \times 10^3$ CFU/mL EAEC isolate EN1E-0227

To test for any adverse effects of *PDX* on the human microbiota, genomic DNA was isolated from triplicate samples of the M10 anaerobic cultures presented in Table 3. High-throughput, multiplex sequencing using Illumina MiSeq, with 150 bp forward and reverse reads were used to analyze DNA from feces only, feces plus *PDX*, feces plus EAEC, and feces plus EAEC and *PDX.* As no DNA was recovered from ciprofloxacin-challenged cultures, no sequence data was generated. After processing and filtering steps, a total of 2303 unique ASVs were detected across all samples.

Alpha diversity (α-diversity), the mean ASV diversity of each sample was calculated using the “estimate richness” function in Phyloseq (Fig. 5). The Shannon diversity index was used to characterize diversity, considering the number of unique taxa (richness) and evenness of taxa present. Mean Shannon indices for Feces only (6.3), Feces + *PDX* (6.3), and Feces + EAEC + *PDX* (6.3) did not differ significantly from one another, but all differed significantly from Feces + EAEC (5.8) which showed the lowest α-diversity (P=0.0035). While the Shannon index utilizes the abundances of each taxa in its diversity calculation, the Chao1 estimator was selected as an additional non-parametric α-diversity estimator for the number of unique taxa present in each sample. Mean Chao1 estimates for Feces only (999.7), Feces + *PDX* (986.7), and Feces + EAEC + *PDX* (1014.2) did not differ significantly from one another, but all differed significantly from Feces + EAEC (682.5), which showed the lowest α-diversity (P=0.0081).

**Figure 5.**
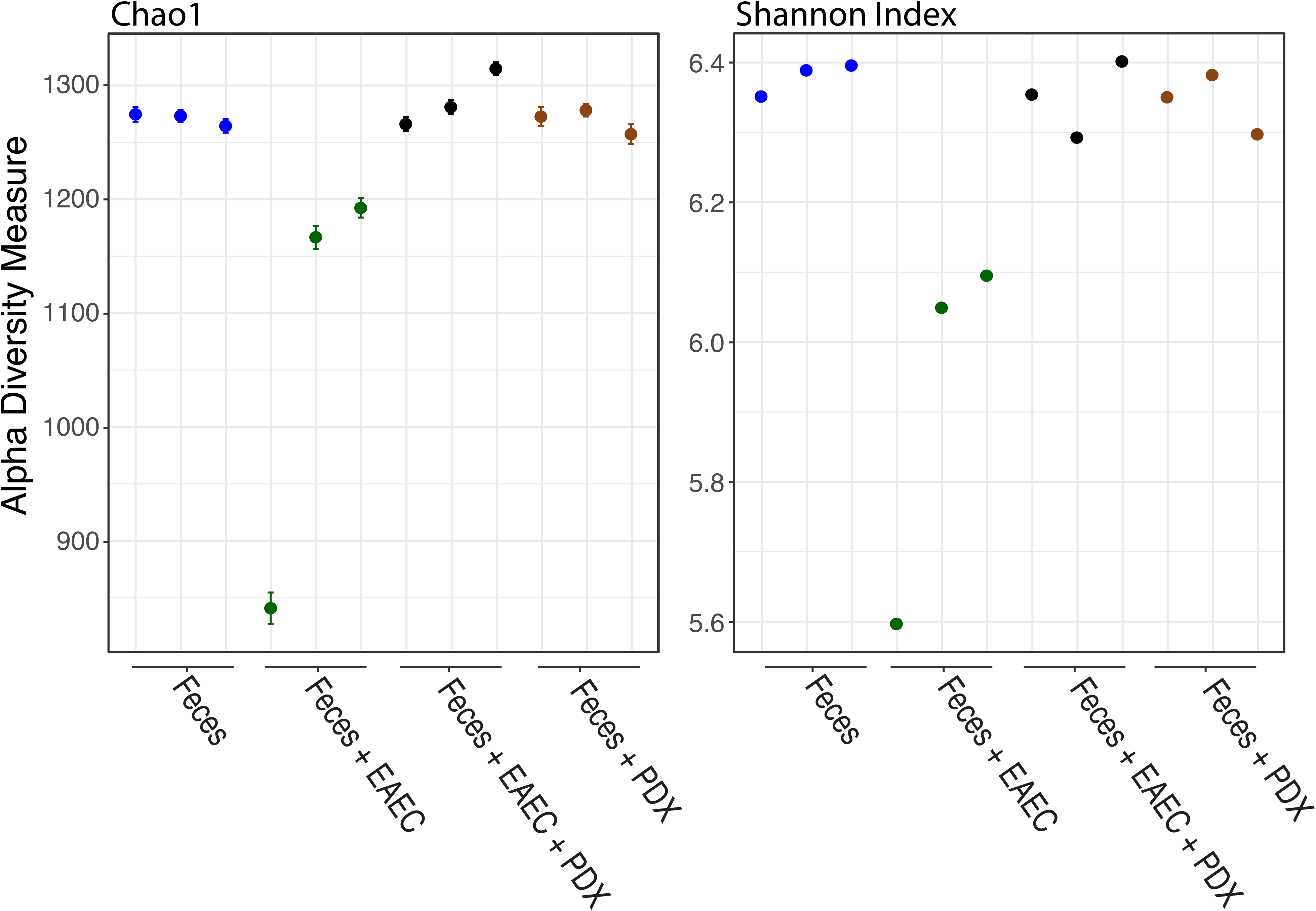
*PDX* does not alter α-diversity of an *in vitro* human microbiome. Chao1 and Shannon indices were used to assess α-diversity of cluster ASV reads. Statistical analyses were carried out to compare mean Chao1 estimates across challenges (n = 12 cultures, Feces only; Feces + EAEC,*, P>0.0081; Feces + PDX; Feces + EAEC + PDX). Statistical analyses were also carried out to compare mean Shannon indices across challenges (n = 12 cultures, Feces only; Feces + EAEC,*, P>0.0035; Feces + PDX; Feces + EAEC + PDX).

While alpha diversity metrics measure microbiome community within samples, beta diversity (β-diversity) is a measure of differences between samples. A weighted-UniFrac dissimilarity matrix was used to plot samples accounting for abundances and phylogenetic relationships of ASVs. Principal coordinate analysis (PCoA) revealed that all samples from Feces only, Feces + *PDX,* and Feces + EAEC + *PDX* cluster closely together, while samples from Feces + EAEC are found far from this cluster (Fig. 6). Using Adonis2 to run PERMANOVA, we find that centroids from treatment groups are significant different from each other (F=2.83, P=0.014), which is primarily driven by the Feces + EAEC group. Dispersion of samples is not significantly different between groups (P=0.647), meeting the assumption of PERMANOVA.

**Figure 6.**
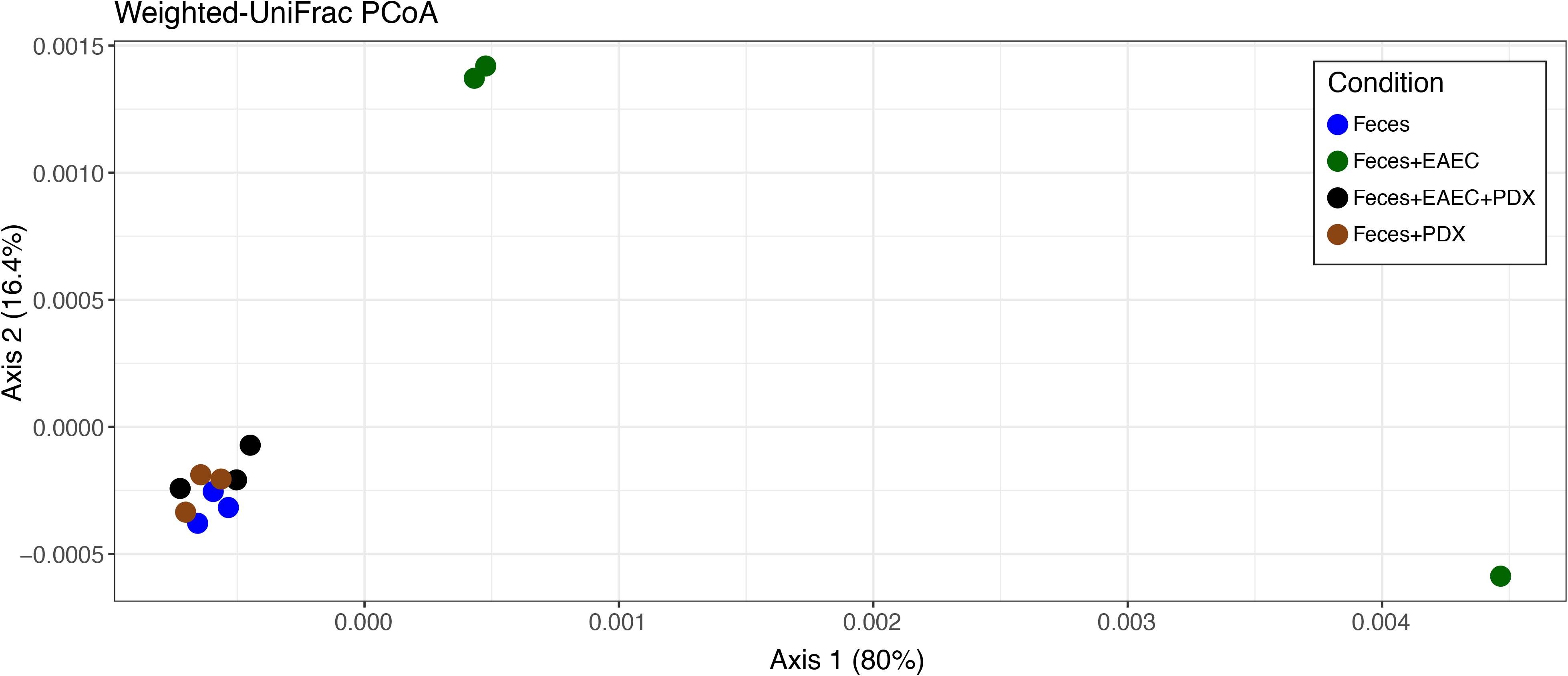
*PDX* does not alter β-diversity of an *in vitro* human microbiome. β-diversity was assessed through principal coordinate analysis (PCoA) for ordination of a weighted-UniFrac distance matrix, accounting for ASV abundance and phylogenetic relationships among taxa. Feces, Feces + *PDX*, and Feces + EAEC + *PDX* conditions cluster together, suggesting microbiome communities are unchanged with the addition of *PDX*, while the addition of EAEC alone alters the community significantly (PERMANOVA: F=2.83, P=0.014)

DESeq2 was used to identify specific taxa that have differential abundances between treatment groups. The only pairwise comparisons that yielded differentially abundant taxa (FDR<0.05) were Feces + EAEC vs. Feces (305 taxa), Feces + EAEC vs. Feces + *PDX* (261 taxa), and Feces + EAEC vs. Feces + EAEC + *PDX* (232 taxa) (Fig. 7). For Feces + EAEC, we observed alteration of the abundance of taxa of all Phyla (except a single representative of the Phylum Verrucomicrobia that was unaffected) in the presence of EAEC isolate EN1E-0227. Taxa of the Bacterioidetes were reduced, as were a number of taxa of genera of the Firmicutes, and Acintobacteria increased compared to the other groups. No comparisons between the Feces, Feces + *PDX*, and Feces + EAEC + *PDX* groups revealed any differentially abundant taxa, even when the FDR cutoff was made less stringent at FDR<0.1.

**Figure 7.**
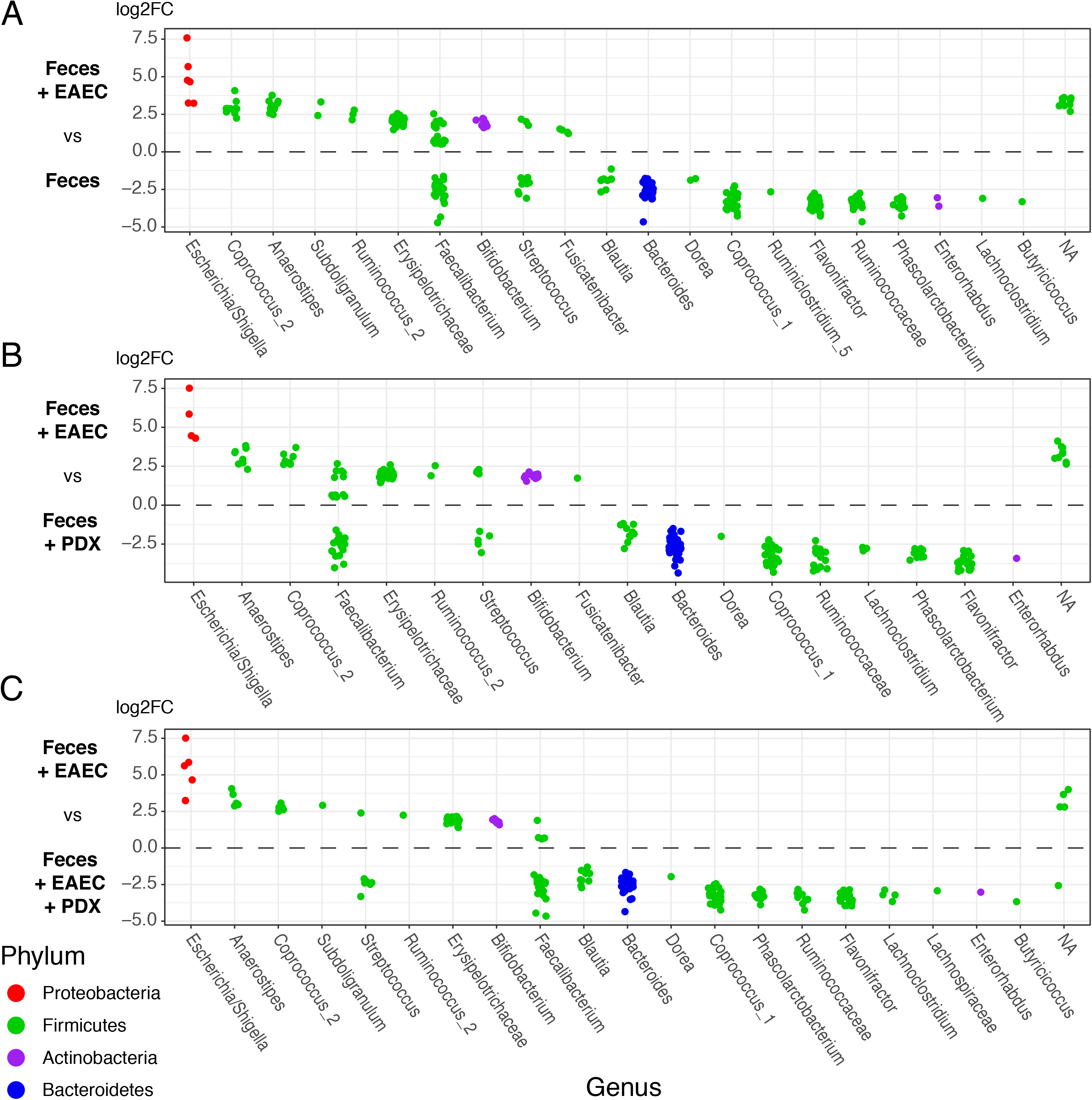
Feces+EAEC is the only condition that shows changes in specific taxa vs controls. Differential abundance analysis was performed via pairwise binomial Wald tests in DESeq2. Among all conditions, comparisons between (A) Feces + EAEC vs. Feces, (B) Feces + EAEC vs. Feces + *PDX*, and (C) Feces + EAEC vs. Feces + EAEC + *PDX* all yielded differentially abundant taxa (P<0.05), while no comparisons among Feces, Feces + *PDX*, and Feces + EAEC + *PDX* showed any significant differences in ASV abundance, suggesting the addition of *PDX* is highly targeted and does not affect the broader microbiome community.

All microbiome analyses were repeated with taxa in the *Escherichia* genus removed to test whether inoculated *E. coli* abundances were driving diversity patterns. However, the results remained largely unchanged for each test. Based on the collective microbiome results, we concluded that *PDX* phage killed EAEC isolate EN1E-0007 when cultured anaerobically in the presence of human gut microbiota without affecting bacterial community diversity.

## Discussion

Using EAEC clinical isolate EN1E-0227 as our target EAEC pathotype, the *Myoviridae Escherichia virus PDX* killed the bacteria both *in vitro* and *in vivo*. EN1E-0227 is a case-associated isolate from a child living in rural Tennessee, and we also demonstrated that *PDX* formed plaques on case-associated EAEC isolates from children in Columbia (Tables 1 and S1). Because of the disparate location of these EAEC isolates, and that *PDX* formed plaques on case-associated EPEC strains from children in the Seattle area, the results suggested that the sequential method (21) for isolating phage that recognized more core bacterial antigens was successful.

In order for phages to be used as therapeutics, they must be strictly lytic, and several lines of evidence indicate that *PDX* is a lytic phage. *PDX* was classified as a member of the strictly lytic *Myoviridae* bacteriophage family (51) based on morphological characteristics from TEM imaging (Fig. 1). Neither CI nor CI-associated genes that are responsible for lysogeny in phage lambda were identified in the genome of *PDX*. It is noteworthy that the genome of *PDX* contains 206 CDS, which is more than the other phages in this cluster in the current Aclame database (Table 2). Analysis of the head-neck structure CDS, in the Virfam suite, identified *PDX* as belonging to the Neck Type One – Cluster 7 (Figs. 2A and B). Importantly, phage induction, by treatment with Mitomycin C, did not induce *PDX* plaques in mutant derivatives of EAEC strain EN1E-0227 resistant to *PDX* infection. We concluded that the *PDX* phage was a member of the *Myoviridae* family having a strictly lytic lifestyle.

Other phages with high sequence identity to *PDX* are being investigate for use as therapeutics or for food contamination interventions. For example, the *Myoviridae* phage Murica (Table S2) is being researched as a potential treatment of contaminated food products (52). *Escherichia* phage V18 was part of a successful prophylactic phage cocktail used in the prevention of traveler’s diarrhea in mice infected with 6 bacterial strains, including *E. coli, Shigella flexneri, Shigella sonnei, Salmonella enetica, Listeria monocytogenes* or *Staphylococcus aureus,* and Lac (-) *E. coli* K-12 C600 (55). Similar to *PDX* replicating in clinical isolates of both EAEC and EPEC pathotypes, the related V18 phage was cited as having a host range that included ETEC, EAEC, and enterohaemorrhagic *E. coli* (EHEC). The ability of *PDX* to kill case-associated strains of two *E. coli* pathotypes, EAEC and EPEC, makes it an ideal candidate for therapeutic administration early, alone or in a cocktail, to patients when a bacterial infection is suspected but the species or strain is unknown, as is common in developing regions of the world. This is particularly the case since EAEC an EPEC bacteria remain a major cause of infantile diarrhea (58). While it is important to use phage cocktails, or multiple phages to combat phage-resistant bacterial mutants (59), in order to develop safe and effective therapies, it is also important to perform *in vitro* and *in vivo* characterization of individual phages (60, 61).

A clear advantage of using lytic phage as therapy against bacterial infections is specific targeting of pathogens without broad destruction of the normal microbiota, though this idea is not established for human infections in the literature. Here we show, by culturing normal human feces anaerobically *in vitro*, that the *PDX* phage kills EAEC EN1E-0227 bacteria in this context, and that by 16S rDNA analysis the α- and β-diversity of the microbiota were unaffected (Figs. 5, 6, and 7). Several points are worth noting. First, by “spiking in” the EAEC strain into the fecal culture, the α-diversity is of course altered (Fig. 5), not unlike when humans are infected with enteric pathogens, and this has also been demonstrated in mice (62). In contrast, the gut microbiome α-diversity was observed to be reduced up to 40% in patients given ciprofloxacin, a fluoroquinolone commonly prescribed for *E. coli* infections (16). There have been many studies that indicate that antibiotic treatment has adverse effects on microbiota diversity (17). In our study, the addition of ciprofloxacin to the anaerobic fecal culture not only killed EAEC strain EN1E-0227, but it obliterated the normal microbiota, such that no DNA was recoverable for the 16S analysis. While a recent study used an *in vitro* human gut culture, chosen to represent a limited bacterial community, to show the phage lyses a target laboratory strain of *E. coli* without significantly altering quantities of mutualistic gut bacteria (63), ours is the first study to our knowledge to address this question with a 16S metagenomic analysis from cultured, normal human feces. The development of antibiotics begins with *in vitro* testing, and thus our approach demonstrates that therapeutic phage can be developed to target pathogens without human gut microbiome dysbiosis.

Dysbiosis from antibiotics leads to loss of keystone taxa, loss of diversity, shifts in metabolic capacity, and blooms of pathogens (16). Similarly, for children living in the developing regions of the world, repeated bouts of diarrhea can lead to dysbiosis and serious health issues (58). These children are at risk for malnutrition, and often suffer from environmental enteropathy: smaller villi, larger crypts, diminished absorption of nutrients and increased inflammation. During bouts of acute diarrhea protective microbiome members, such as Bacterioidetes and Firmicutes, are reduced, creating dysbiosis. In our fecal culture analysis, consistently, in the presence of the EAEC clinical isolate and treatment with phage absent, we observed a reduction in abundance of taxa in the genera belonging to the Bacterioidetes Phylum, as well as reduction in a number of taxa in the genera of the Firmicutes Phylum (Fig. 7). As EAEC is a major contributor to disease in these countries, and is thought to be an underestimated pathogen (1), *PDX* represents a potential therapeutic option, eliminating persistent diarrhea, while helping to support a healthy microbiome to prevent environmental enteropathy in children.

The problem of treating MDR infections will only get more acute and alternate therapy ideas are urgently needed. Phage therapeutics can be one such option. For example, *PDX* and phage with similar characteristics would benefit travelers suffering from MDR, EAEC-mediated diarrhea because it formed plaques on the case-associated, MDR strain BC/Ac338.3 (Table 1). While researchers have discussed the idea of compassionate phage therapy (14), few trials have occurred either in Europe or the US. In the US, phage therapy has mostly occurred for unmet medical needs, as experimental therapies when all other treatment options have been exhausted (https://www.sciencemag.org/news/2019/05/viruses-genetically-engineered-kill-bacteria-rescue-girl-antibiotic-resistant-infection), as phage are able to eliminate bacteria in a manner independent of the MDR phenotype (59, 64). UC San Diego recently launched The Center for Innovative Phage Applications and Therapeutics (IPATH) to refine treatments and to bring phage therapies to market (http://www.sciencemag.org/news/2018/06/can-bacteria-slaying-viruses-defeat-antibiotic-resistant-infections-new-us-clinical). They hope to generate libraries of phages and to use cocktails for individual patients to treat MDR infections. Our efforts and those of other researchers will be essential for isolating therapeutic phages to be used against the increasing number of untreatable bacterial infections, to target specific pathogens but to leave the microbiome healthy and intact.

## Acknowledgments

The authors gratefully acknowledge Dr. Oscar Gómez and Julio Guerra, MS, Division of Pediatrics Infectious Diseases, 875 Ellicott Street, University at Buffalo, The State University of New York University, Buffalo, NY 14203, for providing EAEC strain EN1E-0227 and clinical isolates from children suffering from diarrhea in Columbia.

