## Supplemental Tables S1 and S2 for "*Myoviridae* Phage *PDX* Kills Enteroaggregative *Escherichia coli* without Human Microbiome Dysbiosis"

**Table S1.** EPEC strains[1] used in this study.

| Strain | Serotype | Syndrome[2] | Spot Test Result[3] | Plaque Formation[4] |
|---|---|---|---|---|
| TB85A | 0131:HN | CD | + | - |
| TB96A | 075:HN | BD | + | + |
| TB135A | 0128:H2 | CD | - | - |
| TB156A | O55:H7 | CD | - | - |
| TB171A | O125:H6 | D, V | + | + |
| TB182A | O55:H7 | CD | - | - |
| TB183A | O127:H40 | D | - | - |
| TB204A | O96:HN | D, V, F | + | + |
| TB206A | O26:HN | BD | - | - |
| TB209E | O15:HN | CD | - | - |
| TB216A | O118:H8 | D | - | - |
| TB220B | O153:HN | CD | + | + |
| TB227A | O8:H41 | CD | - | - |
| TB269C | O145:HN | D, F | + | - |
| TB280A | O66:HN | CD | - | - |
| TB309A | O127:H2 | CD | + | - |
| TB320A | ON:HN | BD, V | - | - |
| TB342C | O115:NM | CD | + | - |
| TB353E | ON:HN | CD | + | + |
| TB425B | ON:HN | CD | - | - |

[1] (48)
[2]BD, bloody diarrhea; CD, chronic diarrhea; D, acute diarrhea; F, fever; U, details unknown; V, vomiting.
[3]Plus sign indicates strain is killed by *Escherichia virus PDX.*
[4]Plus sign indicates *PDX* replicates (can form plaques on) the strain.

Table S2. NCBI BLAST[a] results of bacteriophage with the highest whole genome nucleotide percent identity to *PDX*.

| | % Identity with *PDX* Genome[b] | E-Value[c] | Total Scored (max score 255600) [d] | Accession Number |
|---|---|---|---|---|
| *Escherichia* Phage Murica | 97% | 0.0 | 40234 | AKU44119.1 |
| *Escherichia* coli O157 typing phage 4 | 98% | 0.0 | 52649 | AKE45376.1 |
| *Escherichia* Phage JES-2013 | 98% | 0.0 | 52649 | YP_008530272.1 |

*a*: Default BLAST parameters used, with standard 20-amino acid search. *b*: With default parameters, % identity is calculated by subtracting the number of bases and spacers from a perfect identity match. *c:* Statistical test determining the probability that the significance of alignment was due purely to chance. *d:* Under default settings, the BLOSUM62 scoring matrix was used which subtracts 11 points for the existence of a gap with a 1-point deduction for each gap extension made.
